# Likelihood ratio estimation of partial Y-STR profile matches using discrete Laplace models and marginalisation

**DOI:** 10.1101/2025.07.04.663148

**Authors:** Tóra Oluffa Stenberg Olsen, Poul Svante Eriksen, Niels Morling, Mikkel Meyer Andersen

## Abstract

The discrete Laplace model can be used to estimate the evidential weight of a match between the Y-chromosome short tandem repeat (Y-STR) DNA profiles of the evidence material and a suspect. When the weight of evidence of a match between partial Y-STR profiles is assessed, a discrete Laplace model, restricted to the observed loci in the partial evidence profile, is constructed. However, fitting such a discrete Laplace model is time-consuming since it requires estimating the parameters of the discrete Laplace model and validating the model. We implemented the marginalisation method for discrete Laplace models to estimate the evidential weight of a match between a partial Y-STR evidence profile and a Y-STR reference profile. Since the marginalisation method is based on discrete Laplace models for complete Y-STR profiles, no additional models need to be constructed. We compared the likelihood ratios obtained with (1) the marginalisation method and (2) by fitting new discrete Laplace models restricted to the observed loci in the partial evidence profile. In most cases, the two methods yielded similar likelihood ratios.

## 1. Introduction

Y-chromosome short tandem repeat (Y-STR) DNA profiles are widely used in forensic genetic investigations, particularly for analysing DNA mixtures from a female victim and a male perpetrator, in cases, where the male profile can only be determined by analysing Y chromosome markers [1]. The weight of evidence of a Y-STR match between a suspect and the evidence sample can be estimated by a likelihood ratio (LR) [2], i.e.,

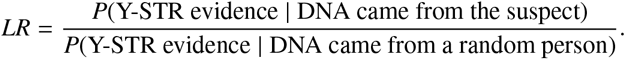

In some cases, the Y-STR evidence profile is incomplete, i.e., one or more loci are missing signals due to, e.g., poor quality of the sample [3]. To assess the evidential weight of partial Y-STR profiles, we assume that the probability of missing a locus does not depend on the allele. Hence, the LR for partial Y-STR profiles reduces to considering only the observed loci. One example, where this assumption is violated, is for degraded samples, where long alleles are more likely to drop-out than short alleles [3, 4].

Statistical models constructed based on databases of complete Y-STR profiles cannot be used directly to assess the evidential weight of partial Y-STR profiles since the profile is not compatible with the model. Hence, the assessment of the weight of evidence needs to be adjusted accordingly.

One approach to estimating the LR of a partial Y-STR profile is to construct a model based on the Y-STR profiles in a reference database restricted to the loci observed in the partial Y-STR profile, i.e., excluding the missing loci of the partial Y-STR profile in all Y-STR profiles in the reference database. Since this model is based on the same loci as the partial Y-STR profile’s, the model can be used directly to estimate the LR of the partial Y-STR profile.

Another approach is to marginalise over the missing loci in the model based on all loci. Hence, no additional model needs to be constructed, and the LR for partial Y-STR profiles is calculated using all complete Y-STR profiles with the partial Y-STR profile as a sub-profile. A more detailed description of this approach for a specific model is given in Appendix A.

We estimated the LR for partial Y-STR profiles using the discrete Laplace method [5] recommended by the International Society for Forensic Genetics [6]. The discrete Laplace method estimates the population frequency of the considered Y-STR profiles to calculate the inverse LR. The discrete Laplace model assumes that the alleles are integers. Thus, intermediate alleles result in partial Y-STR profiles. An expansion of the discrete Laplace model to incorporate such alleles has been proposed [7, 8].

In this paper, we implemented the marginalisation method for the discrete Laplace model to estimate LRs for partial Y-STR profiles. We compared the performance of marginalising over the missing loci in the discrete Laplace model based on all loci to the current approach, i.e., fitting a new discrete Laplace model restricted to the observed loci in the evidence profile. The motivation for proposing a marginalisation in discrete Laplace models is the computational advantages since constructing a new discrete Laplace model is time-consuming, as it requires estimating the parameters of the discrete Laplace model and validating it [9]. Furthermore, the discrete Laplace model depends on the loci of the partial Y-STR profile. Thus, for new partial Y-STR profiles, new models must be constructed.

## 2. Method

### 2.1 Data sets

We considered two data sets of Y-STR profiles. The first data set consisted of 5,804 individuals from Europe [10], and the second data set comprised 1,022 individuals from Denmark (Unpublished). The first data set (EU) is presented in detail in Section 2.1 in [9] as the full reference database *D*^0^ from which we removed Y-STR profiles with null alleles. For the second data set (Denmark), 1,022 male individuals in Danish paternity cases were typed with the AmpFISTR Yfiler^TM^ Plus kit (Thermo Fisher Scientific). The project is registered at the University of Copenhagen’s joint records of processing of personal data in research projects and biobanks (514-0898/23-3000), and it complies with the rules of the General Data Protection Regulation (Regulation (EU) 2016/679). The study follows the policies of the Danish National Center for Ethics (https://nationaltcenterforetik.dk/) and the Danish Research Ethics Committees (https://videnskabsetik.dk/). As in [9], we considered Y-STR profiles of 8, 12, and 17 loci for both data sets. We refer to these sets of Y-STR loci as “kits”, and the particular loci in each “kit” are outlined in [9].

All analyses were done in R [11] version 4.4.1 using the packages disclapmix [12] version 1.7.4.9910, tidyverse [13] version 2.0.0, and ggh4x [14] version 0.2.8. The application code is available in [15].

### 2.2 Reference databases

For the first data set (EU) considered, we drew 15 databases of 550, 1,050, 2,050, and 4,050 Y-STR profiles, respectively. The first 500, 1,000, 2,000, and 4,000 Y-STR profiles, respectively, constituted the databases and the last 50 Y-STR profiles were used for predictions. For the second data set (Denmark), we used all available data and drew 15 reference databases of size 1,022. The last 50 profiles were used for predictions, and the remaining 972 profiles were used for the database. We sampled different Y-STR profiles for predictions each time. We drew 50 × 15 Y-STR profiles without replacement for predictions for the 15 repetitions. For each reference database, the three “kits” introduced in Section 2.1 were considered.

### 2.3 Implementing the marginalisation method for discrete Laplace models

Appendix A shows an expression for estimating the population frequency of a partial Y-STR profile by marginalising over the missing loci in the discrete Laplace model. We showed that marginalising over the missing loci of a partial Y-STR profile resulted in a discrete Laplace model based on only the observed loci of the partial Y-STR profile. Hence, the marginalising in discrete Laplace models is done by using the discrete Laplace model based on complete Y-STR profiles, ignoring the missing loci of the partial Y-STR profile.

We implemented the marginalisation method using the results from Appendix A. The method was added to the disclapmix R package [12] and can be used by specifying “marginalise = TRUE” when estimating population frequencies of partial Y-STR profiles using a discrete Laplace model based on complete Y-STR profiles.

### 2.4 Modelling

For each Y-STR profile used for prediction, we considered the cases where one to five loci were missing in each “kit”. The missing loci were chosen randomly for each Y-STR profile and each “kit”. For each reference database and each “kit”, we fitted the discrete Laplace models *P*full based on the complete reference database and *P*part based on the reference database restricted to the observed loci in the partial Y-STR profile used for prediction.

When fitting the discrete Laplace models, we used the disclapmix adaptive function from the disclapmix package [12], which fits discrete Laplace models with 1, …, *ĉ* + *m* clusters such that the model with *ĉ* clusters yields the lowest marginal Bayesian Information Criterion, BIC, value and where *m* is a user-specific value indicating the number of additional models to test. We chose *m* = 5 as in [9] and the model with *ĉ* clusters. Hence, the choice of *m* = 5 ensures that the marginal BIC could not be improved by examining discrete Laplace models with five clusters more than the chosen model.

### 2.5 Predictions

For each reference database, we extracted the last 50 Y-STR profiles, which were not used when fitting the discrete Laplace models. We estimated the population frequencies of the complete and partial Y-STR profiles by using the discrete Laplace models *P*_ful_l and *P*_part_, and by marginalising using the model *P*_full_ as explained in Section 2.3 and Appendix A. We refer to this “model” as *P*_marg_.

The weight of evidence of a partial Y-STR profile, *x*−*I*, where *I* denotes the set of missing loci in the Y-STR profile *x*, can be estimated as

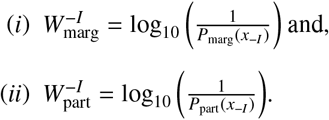

To compare the performance of marginalising in the discrete Laplace model to fitting a new discrete Laplace model, we estimated

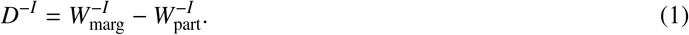

*D*^−*I*^ estimates the increase in the log_10_ of the estimated LR by applying *P*_marg_ instead of *P*_part_.

In addition, when assessing the weight of evidence of partial Y-STR profiles or any subset of complete Y-STR profiles, a consensus has been proposed by [16] for statistical models used to estimate profile or match probabilities stating that: *“The profile (respectively match) probability for the DNA profile of a person on a set of STR loci L is smaller than or equal to the profile (respectively match) probability based on any proper subset of the loci of L”*. In theory, this holds for the discrete Laplace model and, thus, also in practice for the marginalisation method as no parameter is re-estimated. To compare the LR estimates of the complete and partial Y-STR profiles using the discrete Laplace model, where a new discrete Laplace model is constructed to estimate the LR of the partial Y-STR profile, we estimated

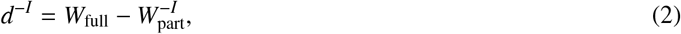

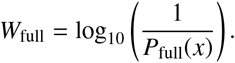

*d*^−*I*^ measures the difference in the log10 of the estimated LR of the complete Y-STR profile compared to the partial Y-STR profile using *P*_par_t. Thus, *d*^−*I*^ ≥ 0 refers to the “cases” where the LR for the complete Y-STR profile was larger than or equal to the LR of the partial Y-STR profile.

## 3. Results

### 3.1 Comparing LRs from the marginalisation method and fitting new discrete Laplace models

Fig. 1 shows the difference in the weight of evidence of marginalising over the missing loci in the discrete Laplace model compared to fitting a new discrete Laplace model restricted to the observed loci of the partial Y-STR profile. Fig. 1 shows quantiles of *D*^−*I*^ defined in Equation (1) for each database size, “kit”, and number of missing loci in the partial Y-STR profile for all “cases”, “cases” where 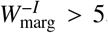5, and “cases” where 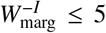, respectively. In particular, the 2.5%, 5%, 50%, 95%, and 97.5% quantiles of *D*^−*I*^ are shown. Fig. 1 shows that the median value of *D*^−*I*^ was approximately zero in all cases. Fig. 1 shows that, for most “cases”, |*D*^−*I*^| < 1. That is, the difference in the weight of evidence obtained by marginalising compared to fitting a new discrete Laplace model was, in most “cases”, less than a factor of ten. Fig. S.1 and S.2 in the Supplementary material show the fraction of “cases” where |*D*^−*I*^| was smaller than a given threshold for each “kit”, database size, and number of missing loci in the partial Y-STR profile for all “cases”, “cases” where 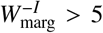, and “cases” where 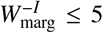, respectively. In addition, Table 1 shows to what extent the LR was increased by at most factors of two and ten, respectively, when marginalising compared to fitting a new discrete Laplace model. Table 1 shows that for 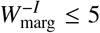, 98 − 100% of the “cases” were increased by less than a factor of ten, while 82 − 94% were increased by less than a factor of two, depending on the “kit” and the number of missing loci. Considering all “cases”, 97 − 100% and 76 − 92% of the “cases” were increased by less than a factor of ten and two, respectively, depending on the “kit” and the number of missing loci.

**Table 1.** Fraction of “cases” below thresholds of *D*^−*I*^, defined in Equation (1), for each “kit” and number of missing loci where we considered all “cases”, “cases” where 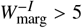, and “cases” where 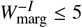, respectively, for the second data set (Denmark). Notice that, no “cases” fulfilled 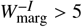 when five loci were missing in the “kit” consisting of 8 loci.

| Number of missing loci | "Kit" | Fraction of "cases" where $D^{-I} < 0.3$ | | | Fraction of "cases" where $D^{-I} < 1$ | | |
| --- | --- | --- | --- | --- | --- | --- | --- |
| | | All | $W_{\text{marg}}^{-I} > 5$ | $W_{\text{marg}}^{-I} \leq 5$ | All | $W_{\text{marg}}^{-I} > 5$ | $W_{\text{marg}}^{-I} \leq 5$ |
| One locus | 8 loci | 0.88 | 0.84 | 0.89 | 0.99 | 0.98 | 1.00 |
|  | 12 loci | 0.83 | 0.79 | 0.89 | 0.98 | 0.97 | 1.00 |
|  | 17 loci | 0.83 | 0.77 | 0.94 | 0.97 | 0.96 | 1.00 |
| Two loci | 8 loci | 0.85 | 0.79 | 0.86 | 0.99 | 0.98 | 1.00 |
|  | 12 loci | 0.84 | 0.79 | 0.87 | 0.98 | 0.95 | 1.00 |
|  | 17 loci | 0.79 | 0.73 | 0.88 | 0.97 | 0.95 | 1.00 |
| Three loci | 8 loci | 0.86 | 0.70 | 0.86 | 1.00 | 1.00 | 1.00 |
|  | 12 loci | 0.84 | 0.76 | 0.87 | 0.99 | 0.99 | 0.99 |
|  | 17 loci | 0.78 | 0.68 | 0.90 | 0.97 | 0.95 | 1.00 |
| Four loci | 8 loci | 0.88 | 1.00 | 0.88 | 1.00 | 1.00 | 1.00 |
|  | 12 loci | 0.87 | 0.73 | 0.90 | 0.99 | 0.99 | 0.99 |
|  | 17 loci | 0.76 | 0.65 | 0.87 | 0.97 | 0.94 | 1.00 |
| Five loci | 8 loci | 0.92 | No "cases" | 0.92 | 1.00 | No "cases" | 1.00 |
|  | 12 loci | 0.84 | 0.71 | 0.85 | 0.99 | 0.96 | 1.00 |
|  | 17 loci | 0.76 | 0.68 | 0.82 | 0.97 | 0.96 | 0.98 |

**Fig. 1.**
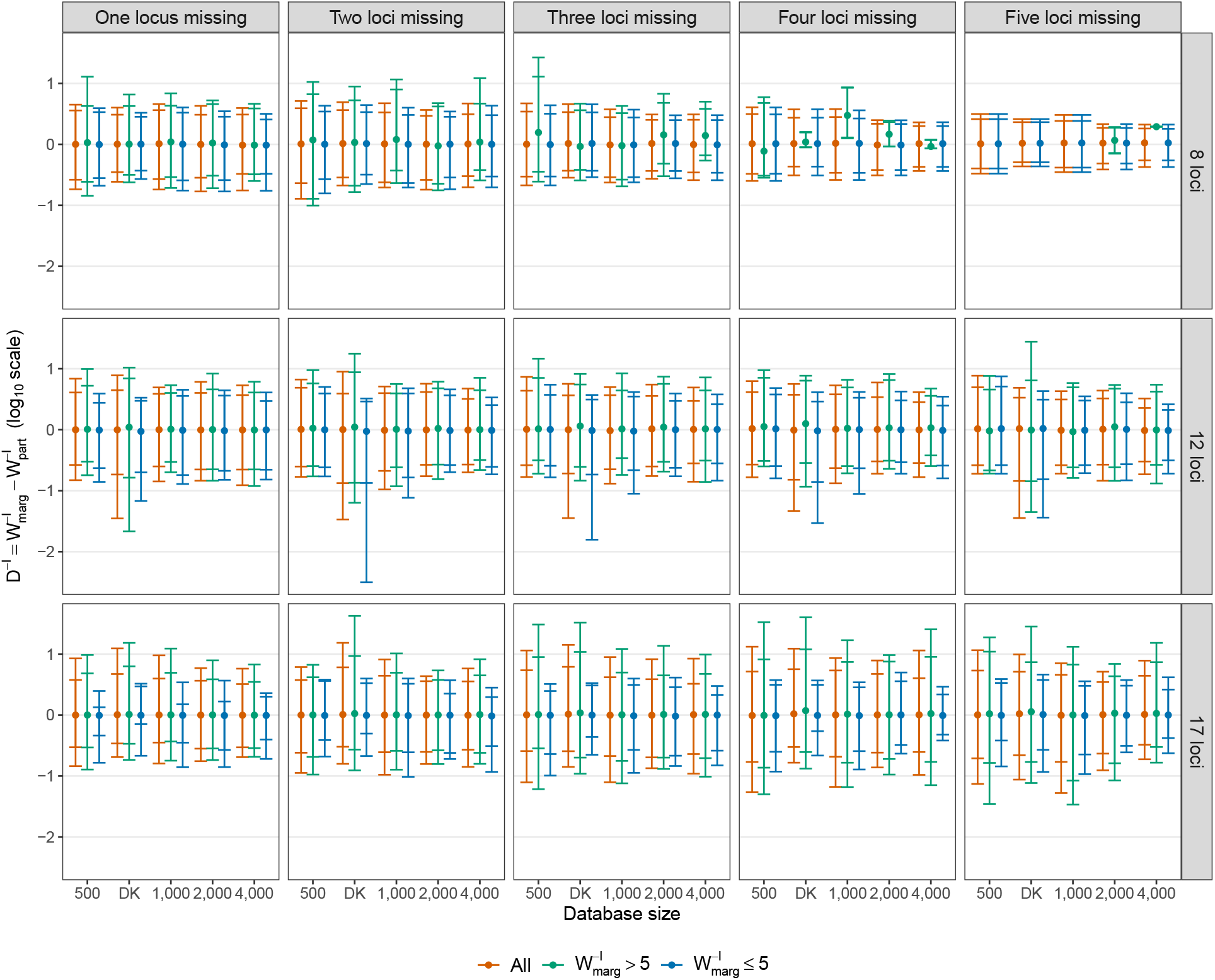
Quantiles of *D*^−*I*^, defined in Equation (1), for each database size and “kit” when one to five loci were missing. That is, e.g., the partial Y-STR profiles used in the top left figure consisted of seven observed loci. The outermost lines represent 2.5% and 97.5% quantiles, i.e., the intervals included 95% of the “cases” considered, and the innermost lines represent the 5% and 95% quantiles, i.e., 90% of the “cases” considered were included in the intervals. In addition, the dots represent the median values of *D*^−*I*^. Notice that “DK” refers to the databases of size 972 extracted from the second data set (Denmark) considered. The groups all “cases”, “cases” where 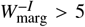, and “cases” where 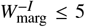 were analysed separately. Notice that, for 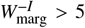 when five loci were missing in the “kit” consisting of 8 loci, intervals are missing since no such “cases” occurred.

### 3.2 Comparing LRs of complete and partial Y-STR profiles using discrete Laplace models

The marginalisation method always results in LRs for partial Y-STR profiles, which are larger than or equal to those of the corresponding complete Y-STR profiles. This follows from the fact that no parameters are re-estimated. Thus, we compare the LR estimates of partial and complete Y-STR profiles when a new discrete Laplace model is fitted to estimate the LR of the partial Y-STR profile.

Fig. 2 shows the log10 of the estimated LR of the complete Y-STR profile plotted against the log10 of the estimated LR of a partial Y-STR profile for each “kit” and each number of missing loci in the partial Y-STR profiles. The black line in Fig. 2 corresponds to the case where *d*^−*I*^ = 0. Fig. 2 is based on only the second data set (Denmark). An analogous figure for the first data set (EU) is shown in Fig. S.3 in the Supplementary material. The majority of the “cases” corresponded to *d*^−*I*^ ≥ 0. However, for some “cases”, *d*^−*I*^ was less than zero, indicating that, in practice, partial Y-STR profiles can result in larger estimated LRs than those of the corresponding complete Y-STR profiles. To quantify the extent to which this occurred, Table S.1 in the Supplementary material shows the fraction of “cases” where *d*^−*I*^ ≥ 0 for each “kit” and number of missing loci. In general, 83 − 100% of the “cases” resulted in *d*^−*I*^ ≥ 0, depending on the “kit” and the number of missing loci. Fig. 2 and Table S.1 show that, in general, the percentages decreased as the number of missing loci increased. For the “kit” consisting of 8 loci, no “cases” resulted in *d*^−*I*^ < 0 when four and five loci were missing, respectively.

**Fig. 2.**
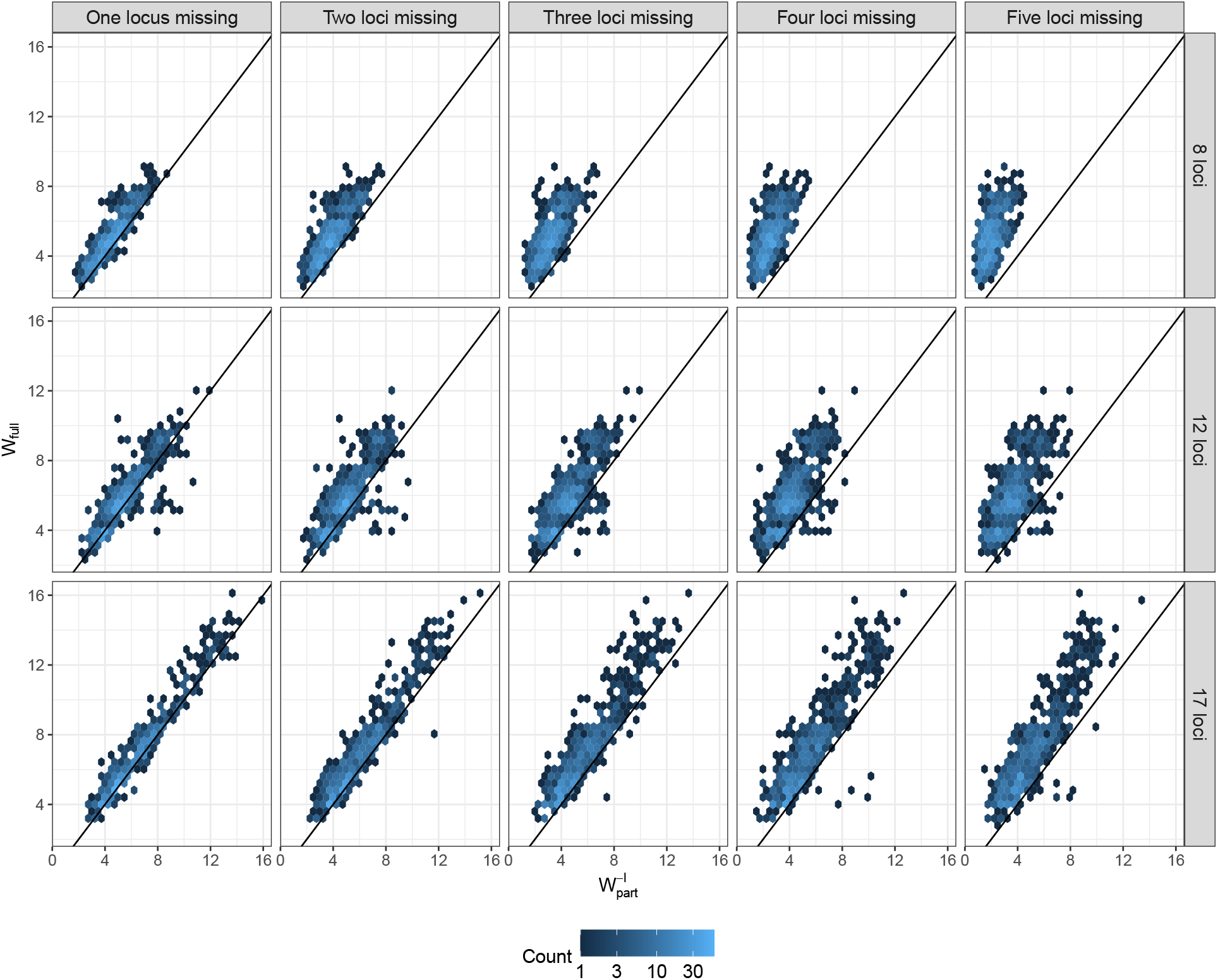
Estimated log10 of the LR of the complete Y-STR profile against the estimated log10 of the LR of a partial Y-STR profile based on constructing a discrete Laplace model restricted to the observed loci of the partial Y-STR profile for each “kit” and number of missing loci in the second data set (Denmark). That is, e.g., the partial Y-STR profiles used in the top left figure consisted of seven observed loci. The black line of slope = 1 and intercept = 0 corresponds to *d*^−*I*^ = 0. The plane is divided into hexagons, where the counts show the number of “cases” in each hexagon. Notice that the total number of “cases” for each figure is 750.

## 4. Discussion

Using Y-STR profiles in forensic genetic investigations involves estimating the weight of evidence of a match between the evidence profile and the Y-STR profile of suspects. We assessed the weight of evidence of partial Y-STR profiles using the discrete Laplace model. To derive an LR for partial Y-STR profiles, we assumed that the probability of missing a locus did not depend on the allele. This assumption can be violated in practice, e.g., with degraded samples. In cases where this assumption is violated, more research is needed to derive an expression for the LR for partial Y-STR profiles.

To estimate the LR for partial Y-STR profiles, the current approach requires fitting a new discrete Laplace model restricted to the observed loci of the partial Y-STR profile. This process can take several minutes to several hours [9], followed by a validation of the constructed discrete Laplace model.

We implemented the marginalisation method for the discrete Laplace model to estimate the weight of evidence for partial Y-STR profiles. We showed that marginalising over the missing loci in the partial Y-STR profile in the discrete Laplace model corresponds to a discrete Laplace model based on only the observed loci in the partial Y-STR profile (Appendix A). Specifically, the method relies on the discrete Laplace model constructed based on complete Y-STR profiles where the missing loci are ignored when estimating the population frequency of the partial Y-STR profile. Hence, no additional discrete Laplace models need to be constructed.

We examined the performance of the marginalisation method, in practice, by comparing the LR estimates to those observed by the current approach, i.e., fitting a new discrete Laplace model restricted to the observed loci of the partial Y-STR profile. We found that, in practice, marginalising in the discrete Laplace model and fitting a new discrete Laplace model restricted to the observed loci gave similar results in the majority of the “cases” considered (Fig. 1 and Table 1). The median difference in the estimated LRs obtained from the two approaches was ca. 0 for all data sets, database sizes, number of loci, and “kits” considered (Fig. 1). Large differences were only observed in a few “cases” (Fig. 1 and S.1 and Table 1). Most “cases” considered resulted in differences between the estimated LRs obtained from the two approaches of at most a factor of two, and almost all “cases” resulted in differences of at most a factor of ten (Fig. 1 and S.1 and Table 1).

We examined the extent to which LR estimates for partial Y-STR profiles were smaller than LR estimates of the complete Y-STR profiles using discrete Laplace models. We found that, in practice, when the population frequency of a partial Y-STR profile was estimated by a discrete Laplace model based on only the observed loci in the partial Y-STR profile on the used data, 83 − 100% of “cases” resulted in LRs of the partial Y-STR profile being smaller than or equal to the LR of the complete Y-STR profile (Fig. 2 and Table S.1). This number increased as the number of missing loci increased (Fig. 2 and Table S.1). Since no parameters are re-estimated when the LR of the partial Y-STR profile is estimated by marginalising over the missing loci in the discrete Laplace model, the LR for the partial Y-STR profile cannot be larger than the LR of the complete Y-STR profile. Based on [16], in general, marginalising to ensure that the LR is not increased for subsets of the complete Y-STR profiles can introduce some issues outlined in the paper. However, since we showed that the two approaches to estimate the evidential weight of the partial Y-STR profiles, i.e., marginalising in a discrete Laplace model and fitting a new discrete Laplace model, yielded similar results in the majority of the “cases”, the presented issues would only occur to a small extent.

## Appendix

**A. Marginalising in discrete Laplace models**

Consider an *l* loci Y-STR profile *x* = (*x*1, …, *xl*). The discrete Laplace model estimates the population frequency of the profile *x* as

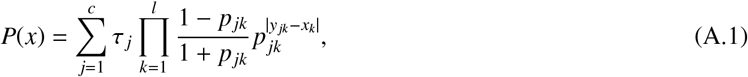

where *c* denotes the number of clusters, τ *j* denotes the probability of a profile belonging to cluster *j, p jk* denotes the dispersion parameter at locus *k* for the *j*’th cluster, and *y jk* denotes the *j*’th centre haplotype at locus *k* [5]. A more detailed description of the method is found in [5].

When marginalising in the discrete Laplace model, the population frequency of a partial Y-STR profile is estimated as a sum of the estimated population frequencies of all complete Y-STR profiles including the partial Y-STR profile as a sub-profile. Thus, if we consider the case where the allele on locus *i* for some *i* ∈ {1, …, *l*} is missing in the Y-STR profile *x* and denote the resulting partial Y-STR profile as *x*−*i* = (*x*1, …, *xi*−1, *xi*+1, …, *xl*), marginalising over this missing allele in the discrete Laplace model yields

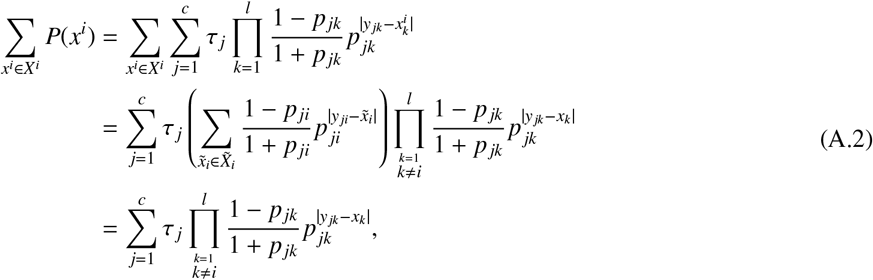

where *X*^*i*^ denotes the set of all Y-STR profiles 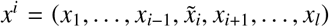 for 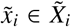 where 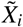 denotes the set of all possible alleles on the locus *i*. Hence, marginalising over the missing allele in the discrete Laplace model based on the complete reference database profiles corresponds to a discrete Laplace model based on only the observed loci in the partial Y-STR profile. Analogously, it can be shown that marginalising over multiple loci results in a discrete Laplace model based on the remaining loci.

