## Supplementary material for "Likelihood ratio estimation of partial Y-STR profile matches using discrete Laplace models and marginalisation"

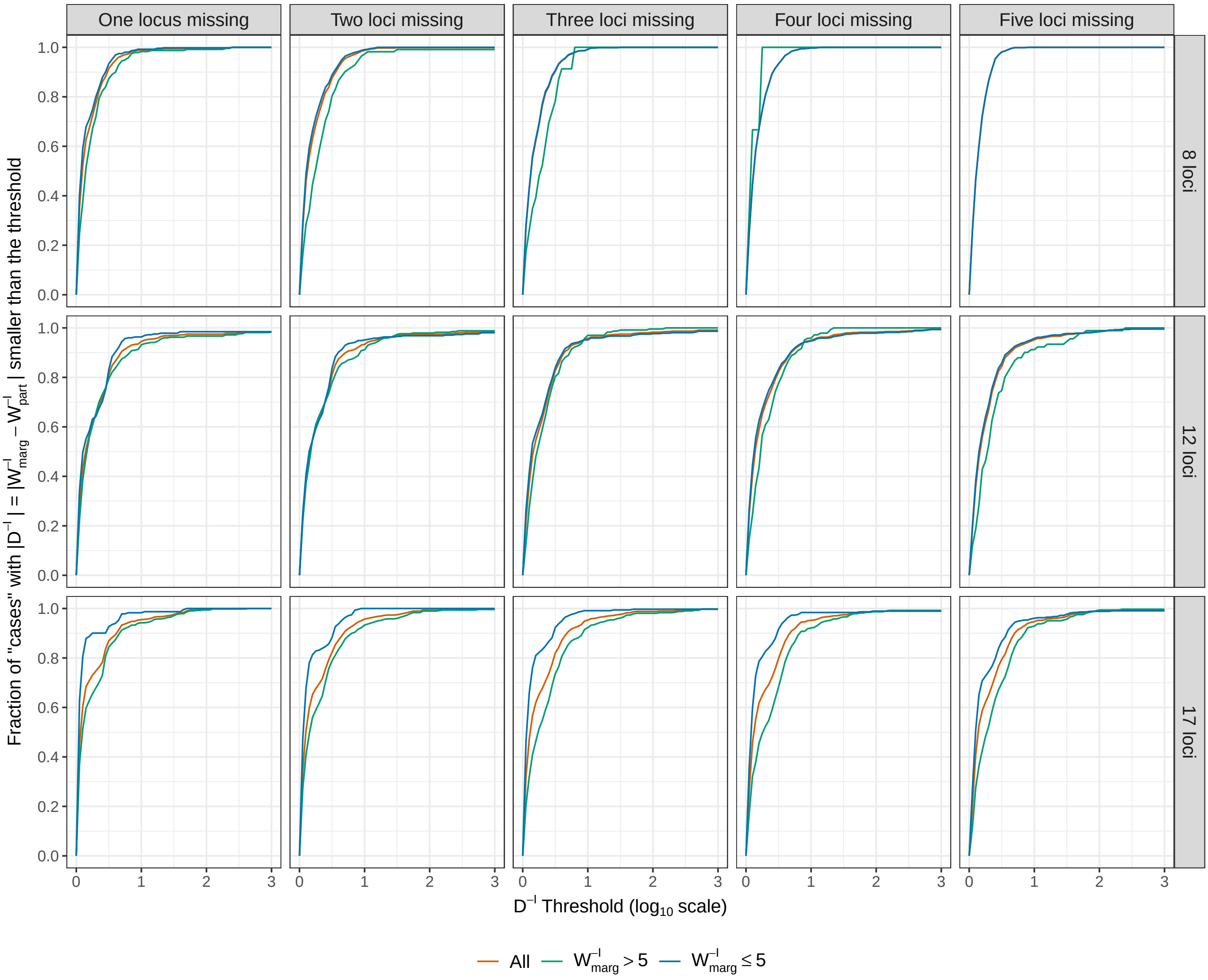


**Fig. S.1:** Fraction of "cases" where $|D^{-I}|$, where $D^{-I}$ is defined in Equation (1), was smaller than a given threshold for each database size and "kit" when one to five loci were missing in the second data set (Denmark). That is, e.g., the partial Y-STR profiles used in the top left figure consisted of seven observed loci. The groups all "cases", "cases" where $W^{-I}_{\text{marg}} > 5$, and "cases" where $W^{-I}_{\text{marg}} \leq 5$ were analysed separately. For example when one locus was missing in the "kit" consisting of 17 loci, ca. 93% of the "cases" resulted in $|D^{-I}| < 0.5$ for $W^{-I}_{\text{marg}} \leq 5$

.

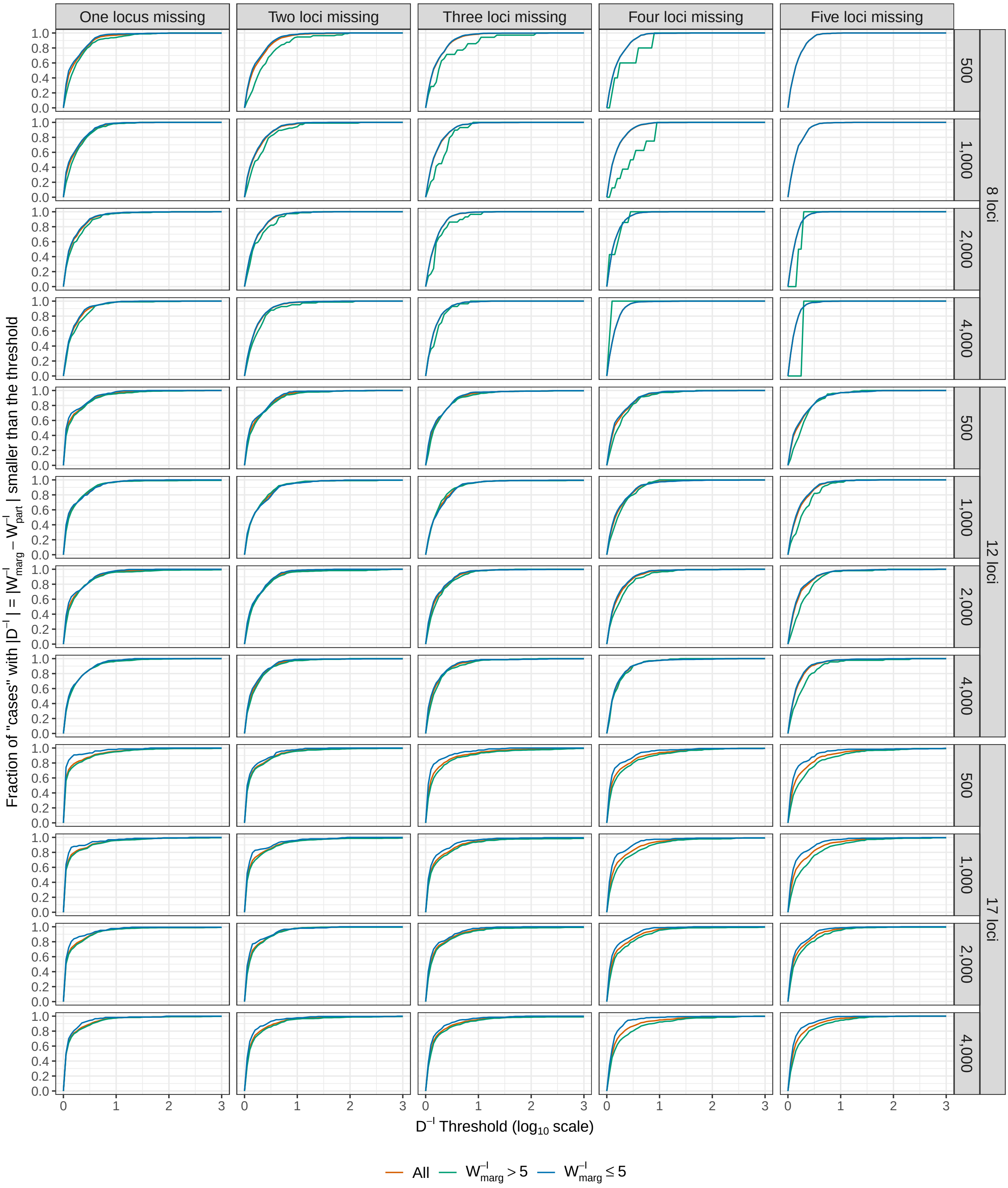


**Fig. S.2:** Fraction of "cases" where $|D^{-I}|$, where $D^{-I}$ is defined in Equation (1), was smaller than a given threshold for each database size and "kit" when one to five loci were missing in the first data set (EU). That is, e.g., the partial Y-STR profiles used in the top left figure consisted of seven observed loci. The groups all "cases", "cases" where $W_{\text{marg}}^{-I} > 5$, and "cases" where $W_{\text{marg}}^{-I} \leq 5$ were analysed separately.

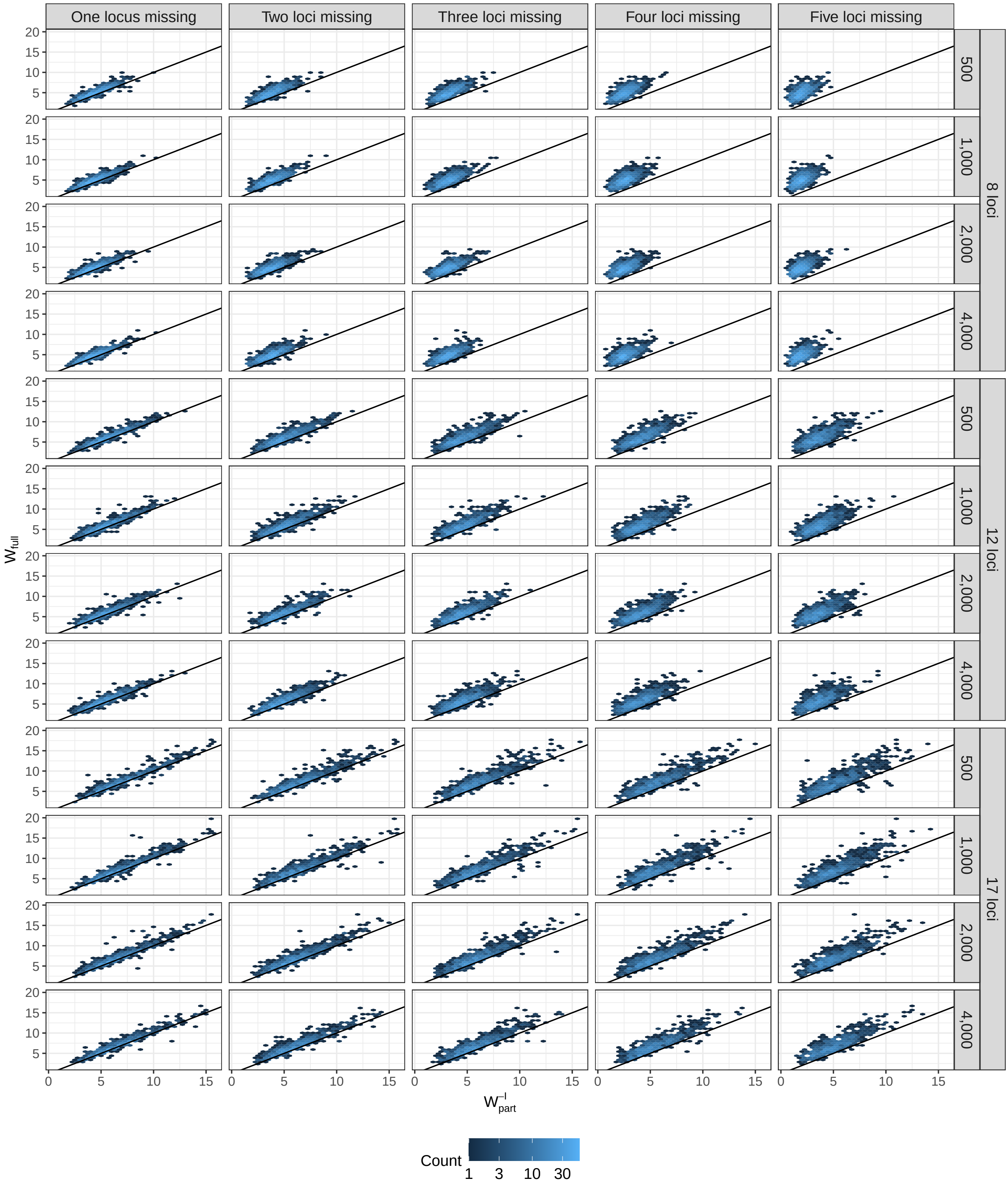


**Fig. S.3:** Estimated $\log_{10}$ of the LR of the complete Y-STR profile against the estimated $\log_{10}$ of the LR of a partial Y-STR profile based on constructing a discrete Laplace model restricted to the observed loci of the partial Y-STR profile for each database size and "kit" when one to five loci were missing in the first data set (EU). That is, e.g., the partial Y-STR profiles used in the top left figure consisted of seven observed loci. The black line of slope = 1 and intercept = 0 corresponds to $d^{-I} = 0$. The plane is divided into hexagons, where the counts show the number of "cases" in each hexagon. Notice that the total number of "cases" for each figure is 750.

**Table S.1:** Fraction of ”cases” where $d^{-I} \geq 0$ for each ”kit” and number of missing loci for the second data set (Denmark).

| Number of missing loci | ”Kit” | Fraction of ”cases” where $d^{-I} \geq 0$ |
|---|---|---|
| One locus | 8 loci | 0.93 |
| | 12 loci | 0.83 |
| | 17 loci | 0.89 |
| Two loci | 8 loci | 0.99 |
| | 12 loci | 0.90 |
| | 17 loci | 0.95 |
| Three loci | 8 loci | 0.99 |
| | 12 loci | 0.93 |
| | 17 loci | 0.97 |
| Four loci | 8 loci | 1.00 |
| | 12 loci | 0.96 |
| | 17 loci | 0.99 |
| Five loci | 8 loci | 1.00 |
| | 12 loci | 0.98 |
| | 17 loci | 0.98 |
